# Integrative analysis allows a global and precise identification of functional miRNA target genes in mESCs

**DOI:** 10.1101/2021.09.24.461622

**Authors:** Moritz Schäfer, Amena Nabih, Daniel Spies, Maxime Bodak, Harry Wischnewski, Patrick Stalder, Richard Patryk Ngondo, Luz Angelica Liechti, Tatjana Sajic, Ruedi Aebersold, David Gatfield, Constance Ciaudo

## Abstract

MicroRNA (miRNA) loaded Argonaute (AGO) complexes regulate gene expression via direct base pairing with their mRNA targets. Current prediction approaches identified that between 20 to 60% of mammalian transcriptomes are regulated by miRNAs, but it remains largely unknown which fraction of these interactions are functional in a specific cellular context. Here, we integrated transcriptome data from a set of miRNA-depleted mouse embryonic stem cell (mESC) lines with published miRNA interaction predictions and AGO-binding profiles. This integrative approach, combined with molecular validation data, identified that only 6% of expressed genes are functionally and directly regulated by miRNAs in mESCs. In addition, analyses of the stem cell-specific miR-290-295 cluster target genes identified TFAP4 as an important transcription factor for early development. The extensive datasets developed in this study will support the development of improved predictive models for miRNA-mRNA functional interactions.

## INTRODUCTION

Mammalian microRNAs (miRNAs) are endogenous small regulatory RNAs, approximately 22 nucleotides in length involved in a diverse array of cellular and physiological processes (Bartel, 2018). They are often found in clusters on the genome and transcribed polycistronically from inter- or intra-genic genomic loci by RNA polymerase II (Lee et al., 2002, 2004). The resulting capped and poly-adenylated primary transcript (pri-miRNA) is processed in the nucleus co-transcriptionally by the Microprocessor Complex which consists of the RNase III DROSHA and the double-stranded RNA-binding protein DGCR8 to produce a miRNA precursor (pre-miRNA) (Denli et al., 2004; Nguyen et al., 2015). After export to the cytoplasm, the miRNA precursor (pre-miRNA) is cleaved by DICER, another RNase III protein (Bernstein et al., 2001; Hutvágner et al., 2001; Zhang et al., 2004), leading to a double-stranded miRNA duplex, which is subsequently loaded into an ARGONAUTE (AGO) protein, the primary effector protein in this RNA interference (RNAi) pathway (Mourelatos et al., 2002). After loading, the AGO-miRNA complexes generally target the 3’ untranslated regions (UTRs) of their target mRNAs at miRNA response elements (MREs) (Lewis et al., 2005), although reports of functional repression via binding to the coding sequence (CDS) exist as well (Hausser et al., 2013; Reczko et al., 2012). MREs are usually complementary to the seed sequence of the miRNA, which is at positions 2-7 of the 5’ end of the mature miRNA (Lewis et al., 2003). MiRNA binding can lead to the functional repression of its target by translation inhibition and decay of the mRNA (Guo et al., 2010).

The rapid rise in discoveries of miRNAs as key regulators of gene expression in many biological processes has prompted an extensive search for their functional targets, as well as the development of tools for a reliable identification thereof. In addition to Watson-Crick base-pairing, other features, such as the evolutionary conservation (Lewis et al., 2005) and the MRE accessibility (Long et al., 2007) have been associated with the regulatory potential of a target site and have been exploited by computational prediction models for the identification of miRNA interactions (Agarwal et al., 2015; Id et al., 2018; Schäfer and Ciaudo, 2020). However, it has been shown that computational models tend to predict a large number of false-positive interactions (Chu et al., 2020; Pinzon et al., 2017). To further improve existing prediction models, the integration of multiple approaches and data sets to obtain more accurate functional miRNA-mRNA interactions has been proposed (Oliveira et al., 2017). These integrative approaches have successfully improved the precision of predictions, but often at the cost of discarding a large number of functional interactions (Davis et al., 2017; Id et al., 2018; Liu and Wang, 2019).

We hypothesize that the inability of current miRNA target prediction models to provide a full and reliable view of functionally relevant miRNA interactions could be partially attributed to the lack of incorporating important context-specific factors. In this study, we performed an integrative analysis combining OMICs data from a unique series of miRNA-deficient mouse Embryonic Stem Cell (mESC) lines generated in the same genetic background (*Drosha*, *Dgcr8*, *Dicer*, *Ago2&1* knockouts) with other publicly available datasets (prediction models, AGO-bound miRNAs) to determine global and accurate direct functional miRNA interactions in mESCs. We further validated our findings by measuring the impact of the deletion of a stem cell-specific miRNA cluster on gene expression. All together we established that only about 6% of expressed genes are subject to direct miRNA-regulation in mESCs, refining previous higher estimations. In addition, we identified an important transcription factor, TFAP4, as a direct miR-290-295 target. TFAP4 is an essential regulator of the Wnt signaling pathway and thus an essential factor for stem cell differentiation.

## RESULTS

### Gene expression is globally perturbed in RNAi knockout mutant mESC lines

To better capture the extent of miRNA-mediated gene regulation in mESCs, we established a unique set of RNA interference knockout (*RNAi*_KO) mESCs using a paired CRISPR/Cas9 approach (Wettstein et al., 2016) (*Dgcr8_*KO (Cirera-Salinas et al., 2017), *Drosha*_KO (Cirera-Salinas et al., 2017), *Dicer*_KO (Bodak et al., 2017a) and *Ago2&1*_KO (Figure S1A)). For each *RNAi*_KO line, two independent clones were generated, and the downstream protein depletion was validated by Western Blotting (WB) (Figure S1B). We subsequently performed transcriptome profiling of small RNAs (sRNAs) and messenger RNAs (mRNAs) in duplicates from wildtypes (WT) and each *RNAi*_KO line. PCA plots for both RNA-seq experiments generally showed clustering of biological replicates and separation of samples (Figure 1A, Table S1 RNA-seq and Figure S1C, Table S2 sRNA-seq). In the RNA-seq data, *Dgcr8*_KO replicates showed minor differences in gene expression levels and, unsurprisingly, clustered with *Drosha_*KO samples, its Microprocessor complex partner. Additionally, as expected, the *RNAi*_KO mESC lines *Dgcr8*_KO, *Drosha*_KO and *Dicer*_KO exhibited decreased miRNA expression levels (Figure S1D). Interestingly, the depletion of both Argonautes also led to a strong reduction of the vast majority of miRNAs, suggesting that miRNAs are degraded in the absence of the AGO1 and AGO2, which are the main Argonaute proteins expressed in this cell type (Müller et al., 2020).

**Figure 1:**
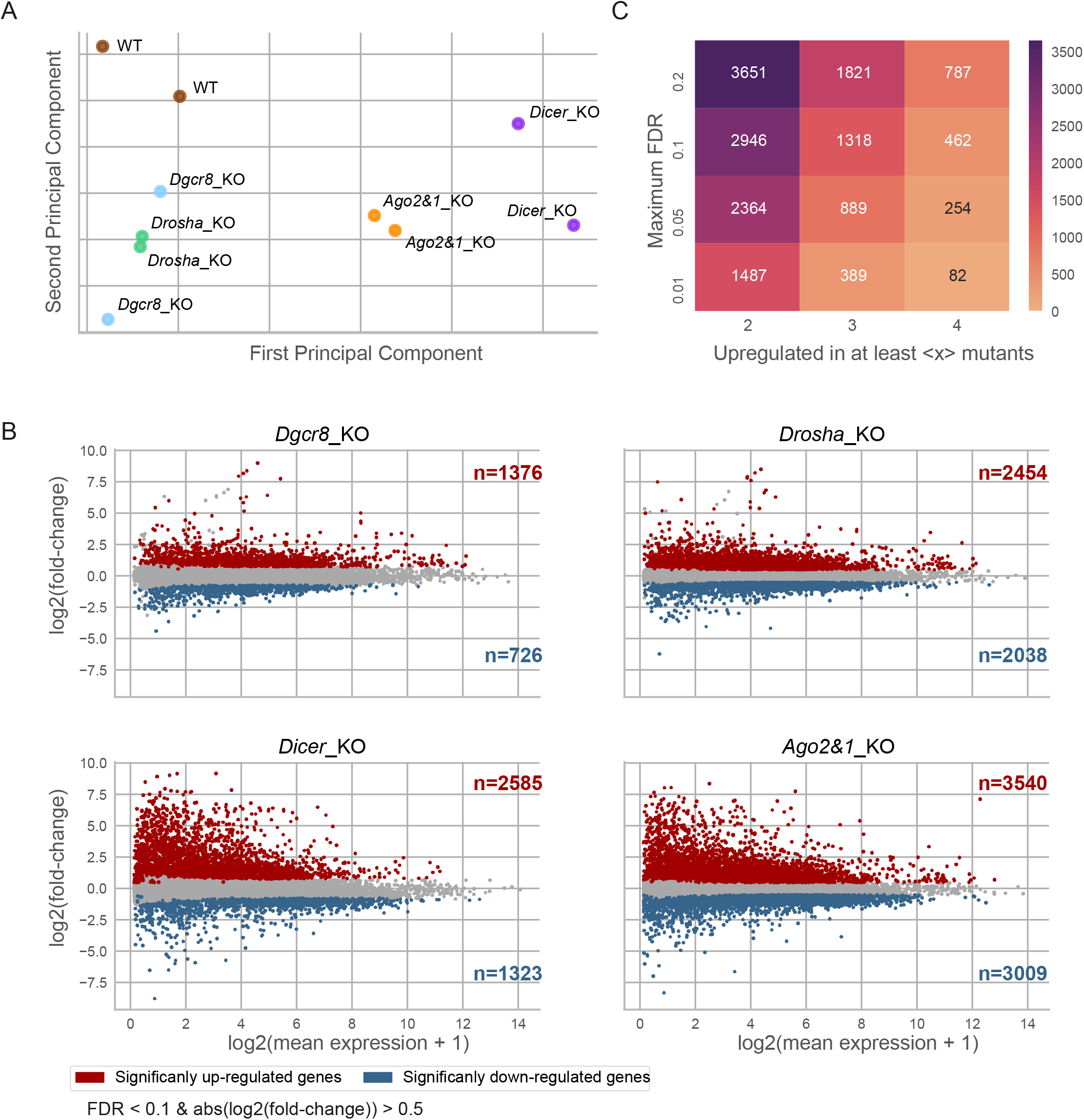
Gene expression is globally perturbed in *RNAi*_KO mESCs. (A) PCA plot of gene expression as measured by RNA sequencing (RNA-seq) in *RNAi*_KO and WT samples. Biological replicates are indicated with the same color. (B) MA plots of the DGE analysis in *RNAi*_KO mutants. Significant up- and downregulated genes are determined based on cutoffs (FDR < 0.1 and log2(fold-change) > 0.5 / < −0.5) and colored in red and blue respectively. (C) Heatmap indicating the number of genes commonly upregulated across mutants for two different selection dimensions: Y-axis denotes maximum false discovery rate (FDR) to consider a gene as *upregulated*. Matrix fields correspond to the number of *upregulated* genes, identified in at least 2, 3 or 4 mutants, as denoted by the x-axis.

To assess the effect of miRNA depletion on gene expression, we performed a differential expression (DE) analysis on the RNA-seq data. We expected the loss of miRNA-mediated regulation to lead to an upregulation of their target genes. Indeed, we observed a large number of upregulated genes in all *RNAi*_KO mutants (from 1376 in *Dgcr8*_KO to 3540 in *Ago2&1*_KO cell lines) (Figure 1B). However, the overlap across all mutants was unexpectedly small (462 genes, false discovery rate (FDR) < 0.1, Figure 1C). In addition, the Pearson correlation for differential gene expression (DGE) values of *RNAi*_KO cell lines was positive for all sample pairs; the correlation coefficients were high for Drosha/Dgcr8 and Dicer/Ago2&1 pairs (0.81 and 0.8 respectively) but rather low (0.34 – 0.53) for all other pairs (Figure S1E), pointing to differences between nuclear and cytoplasmic processes across *RNAi*_KO mESC lines.

Finally, all mutants exhibited a sizeable number of downregulated genes (Figure 1B), with only 126 overlapping between all genotypes (FDR< 0.1, Figure S1F), most likely indicative of numerous indirect effects of the primary loss in RNAi factors.

In conclusion, the extended perturbation of gene expression observed in *RNAi*_KO mESC lines is not exclusively caused by the lack of miRNAs. Mutant-specific effects and secondary regulation events seem to strongly influence global gene expression profiles in stem cells.

### Integrative transcriptomics data analysis identifies global maps of functional miRNA interactions

To identify direct and functional miRNA-mRNA pairs with high accuracy, we integrated sRNA-seq and RNA-seq data with published AGO2 binding data (Li et al., 2020) and, sequence-based interaction prediction data from TargetScan (Agarwal et al., 2015) as visualized in Figure 2A. Briefly, potential interactions were established based on seed matching at AGO2 binding peaks and then filtered based on WT miRNA expression and target mRNA upregulation in *RNAi*_KO mESC lines. Finally, the sum of all four normalized features led to an interaction score, which was used to rank interactions and genes based on the evidence supporting their of regulation by miRNAs.

**Figure 2:**
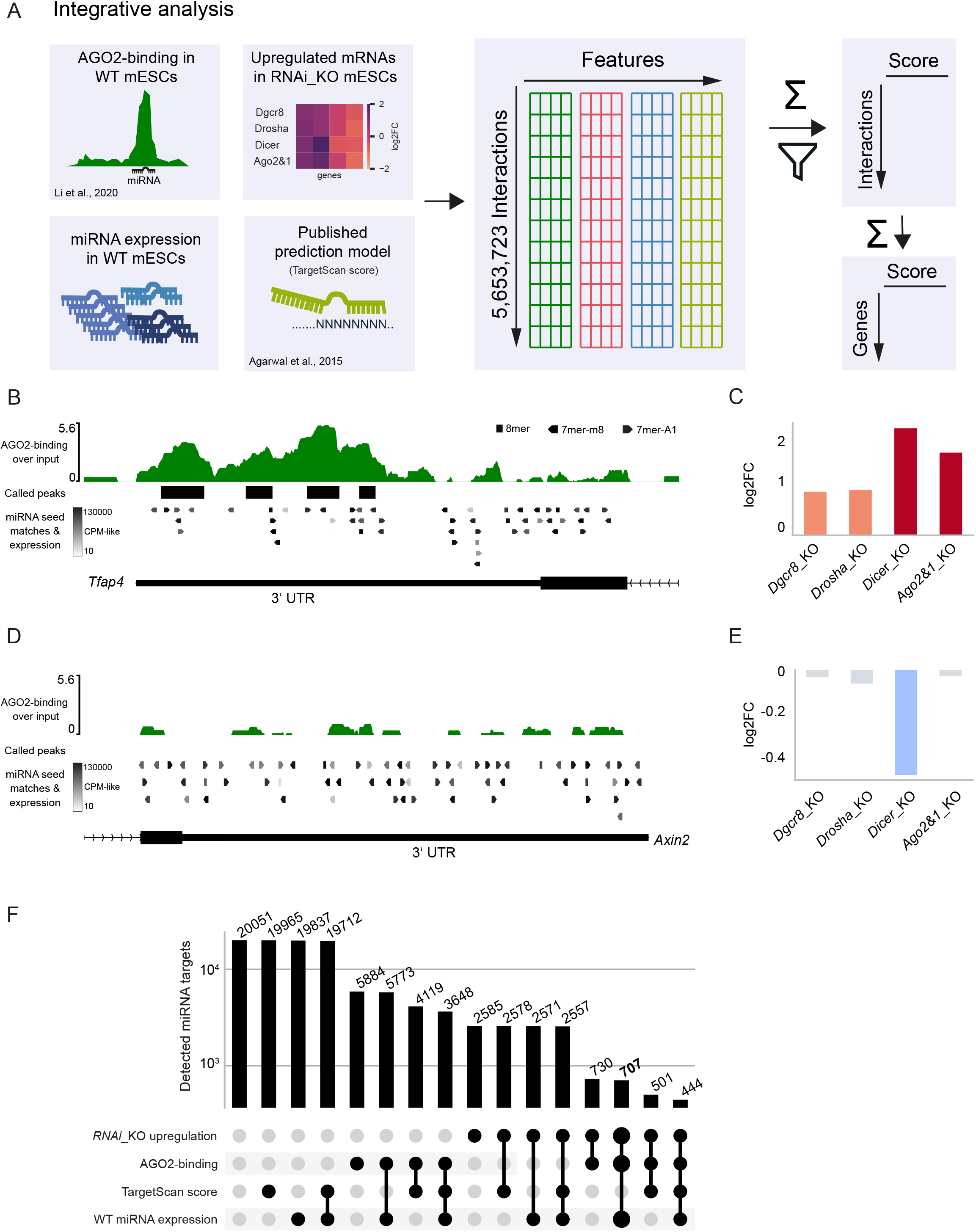
Multi-OMICs integration allows the identification of functional miRNA interactions in mESCs. (A) Graphical overview of data sources and integration. Data are integrated on a per-interaction basis, filtered by miRNA expression, AGO2-binding, target upregulation and scored on a per-interaction and per-gene basis. Filtering and scoring are described in detail in the Materials and Methods section. The latter allows for a certainty-ranking of interactions and target genes. (B and C) Example of integrated data for the *Tfap4* gene with multiple evidences for functional miRNA interactions. (B) shows AGO2-binding profile and called peaks as obtained from (Li et al., 2020), along with predicted miRNA binding sites with the expression of the corresponding miRNA in WT mESCs (Only binding sites for miRNAs with minimal expression of 10 *CPM-like* are shown). (C) shows the misregulation of *Tfap4* in *RNAi*_KO mutants in log2FoldChanges compared to WT. (D and E) Example of integrated data for the *Axin2* gene with no evidence for functional interactions. Same data are shown as in (B, C). (F) Number of identified functional miRNA target genes for different integrative filtering approaches. The integration and filtering by mutant upregulation and AGO2-binding data leads to a restrictive selection of target genes. Reported targets (the bold column) were not filtered by TargetScan scores to avoid filtering of interactions in the coding sequence (CDS) and 5’UTR.

We identified high confidence miRNA-mRNA pairs as, for example, shown in Figures 2B, C, while discarding putative interactions with little evidence as, for example, shown in Figures 2D, E and S2A-D. Filtering interactions by all four integrated features narrowed the number of detected miRNA target genes down to 444 (Figure 2F, Table S3). Consistent with other studies (Hausser et al., 2013; Patel et al., 2020; Reczko et al., 2012), we observed seemingly functional interactions in regions outside of the 3’UTR (for example Figures S2E, F), which led us to exclude the 3’UTR-centric TargetScan context++ score model from the filtering, still leading to a surprisingly low number of 707 identified miRNA-targeted genes (Figure 2F).

In conclusion, multi-OMICs integration facilitated the exclusion of nonfunctional miRNA-mRNA interactions to obtain a set of high-confidence miRNA target genes in mESCs.

### Ribosome profiling in *RNAi*_KO mESCs reinforces the identification of functional miRNA target genes

A substantial part of the data used in the integrative analysis was based on transcriptomics approaches, and we wondered whether the upregulation of miRNA targets in *RNAi*_KO mutants was also reflected at the translational level. We thus assessed the full proteome of all RNAi mutant and WT mESCs using sequential window acquisition of all theoretical fragment ion spectra mass spectrometry (SWATH-MS) experiments (Gillet et al., 2012), and measured the differential protein abundance of predicted miRNA targets. This approach allowed us to capture protein abundances only for 26% of predicted miRNA targets (186 of 707, Figure S3A). All *RNAi*_KO mutants showed significantly enriched positive log2FoldChanges (log2FCs) as compared to a control distribution of log2FCs (p<0.002 for every mutant, student’s t-test), with at least 60% of genes exhibiting positive log2FCs (Figure S3A, Table S4). Since the experiment only captured about a quarter of predicted target genes, we next performed Ribosome profiling (Ribo-seq) in *RNAi*_KO mESCs to measure the ribosome-occupancy on mRNAs (Brar and Weissman, 2015). The high sensitivity of the approach allowed us to detect and compare ribosome occupancy for 95% of predicted miRNA targets (670 of 707, Figure 3A). For every mutant, enrichments for positive log2FCs were statistically more significant and stronger than in the mass spectrometry experiment (p<2e-13 for every mutant, Figure 3A, Table S5). *Drosha*_KO, *Dicer*_KO and *Ago2&1*_KO showed the strongest enrichment (77%, 74%, 77% positive log2FCs respectively), while the enrichment in *Dgcr8*_KO was a little weaker (64% positive log2FCs).

**Figure 3:**
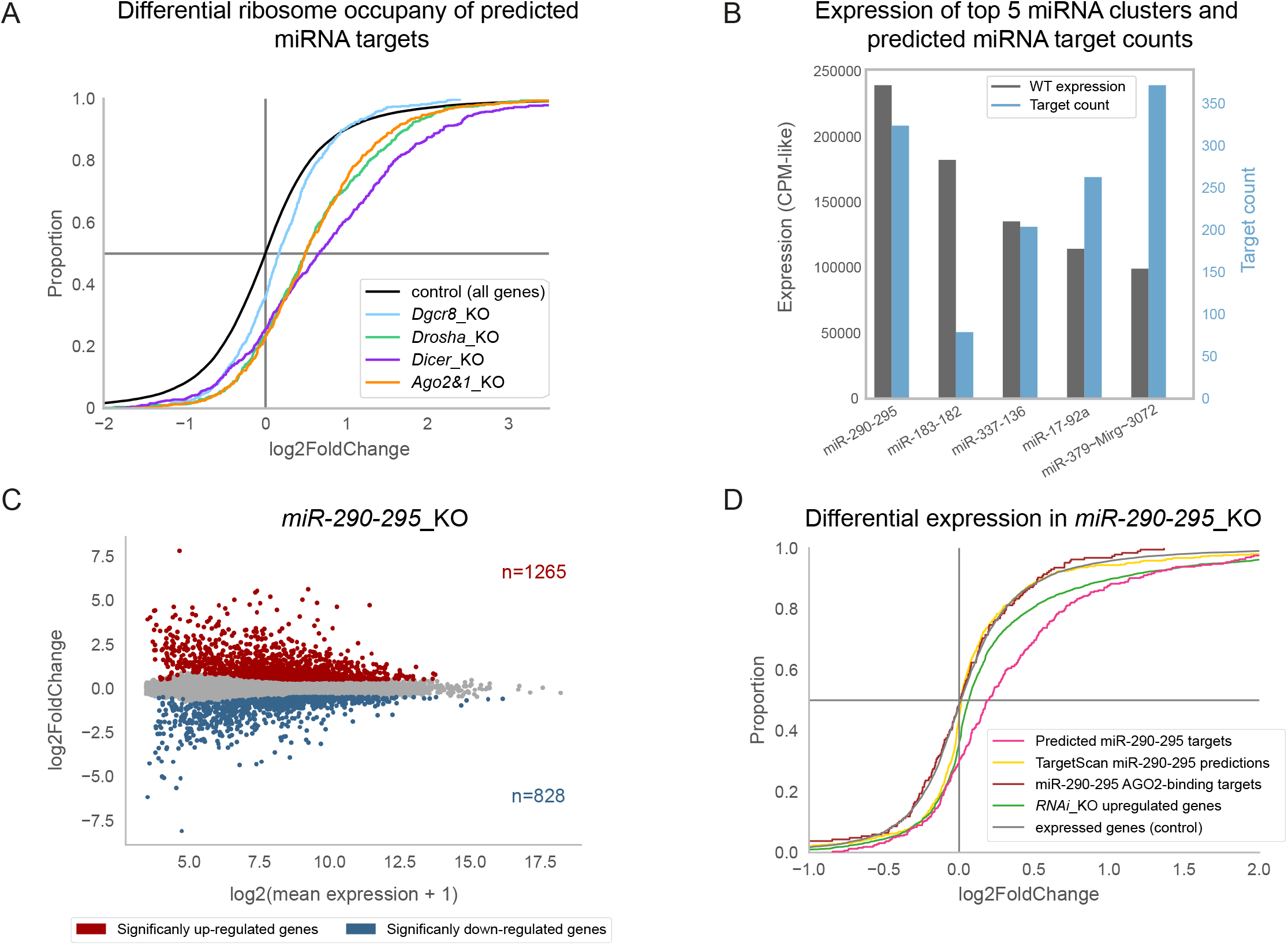
Validation and refinement of miRNA interaction predictions. (A) Cumulative distribution function of differential ribosome occupancy as detected by ribosome profiling (Ribo-seq) in *RNAi*_KO mutants. Of 707 predicted miRNA target genes, 670 (95%) were detectable in the Ribo-seq data and appear in the plot. (B) Bar plot of expression and number of predicted target counts for top 5 expressed miRNA clusters. (C) MA plot of the DGE analysis in *miR-290-295*_KO mESCs. Significant genes were selected using thresholds of FDR < 0.1 and log2FoldChange > 0.5 / < −0.5. (D) Cumulative distribution function of differential expression in *miR-290-295*_KO for different gene groups. Colored gene groups correspond to different identification methods for miR-290-295 target genes based on different data sets. The pink line/group corresponds to 324 miR-290-295-targeted genes out of the 707 miRNA target genes that have been identified in the integrative analysis (Figure 2A).

Taken together, the Ribo-seq approach reinforces the validity of our integrative analysis and the list of functional high-confidence miRNA targeted genes in mESCs.

### Combination of DEG from several *RNAi*_KO mESC lines is a key feature for the identification of functional miRNA targets

Mouse ESCs have a specific miRNA expression pattern (Ciaudo et al., 2009; Houbaviy et al., 2003), with over 50% of all expressed miRNAs originating from four genomic clusters (Calabrese et al., 2007). Indeed, the miR-290-295 cluster is the most strongly expressed cluster in our dataset and almost half of our predicted miRNA target genes are predicted to be targeted by at least one member of this cluster (324 out of 707 genes, Figure 3B). MiR-290-295 has previously been shown to be involved in pluripotency maintenance and self-renewal of mESCs through several molecular processes (Yuan et al., 2017). We therefore complemented our predictions with experimental data by deleting the miR-290-295 cluster in the same mESC background as the *RNAi*_KO mESCs, using a similar paired CRISPR/Cas9 approach (Figure S3B) (Wettstein et al., 2016). We generated two independent cell lines and confirmed the integrity of the deletion at DNA and RNA levels (Figures S3C, D). We next profiled the transcriptome of the *miR-290-295*_KO mESCs and performed a DE analysis (Table S6). Surprisingly, *miR-290-295*_KO mESCs exhibited a strongly misregulated transcriptome with 1265 up- and 828 downregulated genes, similar to the *Dgcr8*_KO mESC lines (Figures 1B, 3C). We then specifically looked at the DE of predicted miR-290-295 target genes in the *miR-290-295*_KO mESC lines. As expected, predictions from the integrative analysis (324 predicted miR-290-295 targets) showed a strong enrichment for upregulation, with 70% of predicted genes (226 of 324) exhibiting a positive log2FC (pink curve in Figure 3D, Table S6). We used the DE analysis to estimate the predictive contribution of each individual dataset used in our integrative analysis. As shown in Figure 3D, predictions based on AGO2-binding and on TargetScan scores showed virtually no enrichment for positive log2FCs, while predictions based on *RNAi*_KO upregulation showed a minor enrichment for upregulated genes, however to a lower degree than the integrative analysis (60% of targets showed a positive log2FC, Figure 3D).

These results demonstrate that the DE across several *RNAi*_KO cell lines is an important feature for the prediction of functional miRNA targets. Further, generation of the *miR-290-295*_KO mutant cell lines allowed for the identification of a high-confidence set of miR-290-295 target genes.

### Depletion of the miR-290-295 cluster combined with predicted functional interactions identifies novel key transcription factors regulated by miRNAs

Out of the 324 predicted miR-290-295 targets, 92 showed a statistically significant upregulation in the *miR-290-295*_KO mutant (Table S6, Figure S4A). Interestingly, 13 of these were annotated as transcription factors (TFs) by the mTFkb database (Sun et al., 2017) of which three of them, *Tfap4, Dazap2 and Mycn,* had previously been implicated in stem cell functions including pluripotency (Chappell and Dalton, 2013; Papathanasiou et al., 2021; Sugawara et al., 2020) (Figure S4A). TFs are proteins that primarily bind to promoter regions to modulate gene transcription (Spitz and Furlong, 2012), therefore potentially contributing to the observed DGE that cannot be explained by the depletion of miRNAs alone. *Tfap4* showed the highest interaction score from our analysis (Figures S4A,B) and has recently been shown to be required for reprogramming mouse fibroblasts into pluripotent stem cells (Papathanasiou et al., 2021). We first investigated its regulation by miRNAs in mESCs by monitoring its expression at the protein level in *RNAi*_KO mutants (*Drosha*_KO and *Dicer*_KO) and *miR-290-295*_KO mESC lines. We observed an upregulation of at least 2-fold for TFAP4 in all mutant cell lines (Figure 4A). To assess whether this upregulation was caused by miR-290-295-mediated repression, we transfected *miR-290-295*_KO mESCs with two miRNAs of the miR-290-295 cluster (miR-291a-5p and miR-291a-3p) and measured TFAP4 expression by WB. Indeed, we observed a downregulation of TFAP4 protein levels as a result of miR-291a-5p and miR-291a-3p transfection (Figure 4B) and an even stronger downregulation of TFAP4 upon transfection with the two miRNA mimics simultaneously. These data strongly suggest that *Tfap4* is regulated by these two members of the miR-290-295 cluster.

**Figure 4:**
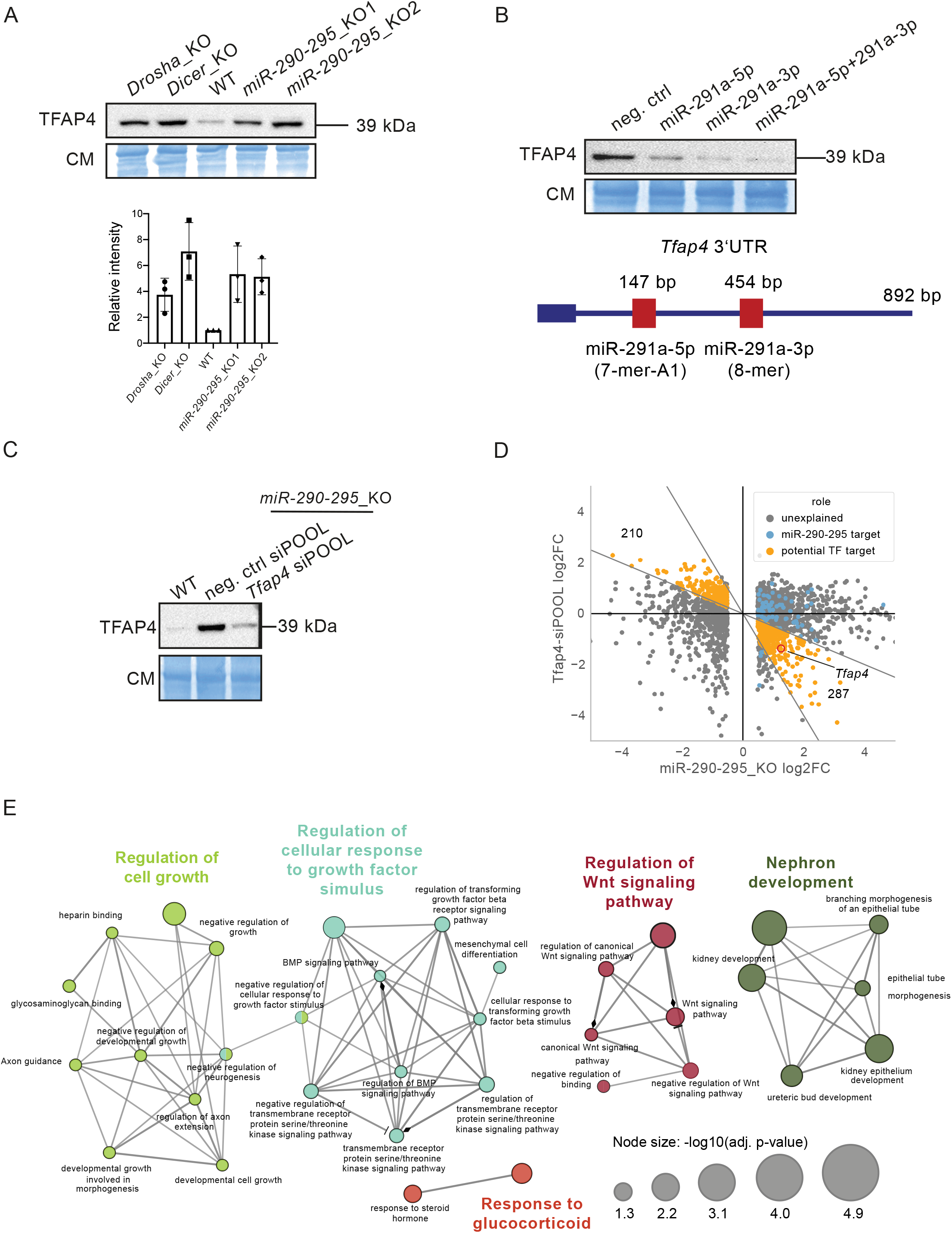
*Tfap4* is a key regulator of gene expression in mESCs. (A) Top: Immunoblot analysis of TFAP4 in *Drosha*_KO, *Dicer*_KO, Wild type (WT), *miR-290-295*_KO1 and *miR-290-295*_KO2 mESCs. Immunoblots were stained with Coomassie blue dye as loading control. Blot is a representative image of three biological replicates. Bottom: Bar graph showing quantification of TFAP4 intensity, normalized to Coomassie and relative to the WT sample in three biological replicates. (B) Top: *miR-291a* regulates TFAP4 expression in mESCs. Immunoblot analysis of TFAP4 after transfection of miRNA mimics (miR-291a-5p, miR-291a-3p and miR-291a-5p+miR-291a-3p combined) in *miR-290-295*_KO2 mESCs. Immunoblots were stained with Coomassie blue dye as a loading control. Blot is a representative image of three biological replicates. Bottom: Map of *Tfap4* 3’UTR indicating miRNA response elements for the transfected miRNA mimics. (C) Immunoblot validation of siPOOL-mediated knock down of TFAP4. TFAP4 levels were compared between untreated WT versus *miR-290-295*_KO2 cells treated with a negative control and a *Tfap4*-targeted siPOOL. Immunoblots were stained with Coomassie blue dye as a loading control. Blot is a representative image of two biological replicates. (D) Scatterplot of differential gene expression in *miR-290-295*_KO control versus siPOOL-*Tfap4*. SiPOOL experiments were performed in *miR-290-295*_KO2 cells and DE was assessed relative to a negative control siPOOL transfection. Only genes that are statistically significantly differentially expressed in *miR-290-295*_KO2 are shown. Genes predicted to be targeted by miR-290-295 are marked in blue. Genes are defined as rescued (orange) if the log2FoldChange-ratio between *miR-290-295*_KO control and siPOOL-*Tfap4* is in the range [−2, −1/2]. *Tfap4* is marked by a red circle. (E) Gene ontology analysis of 121 putative *Tfap4* target genes using ClueGO. Only members of the top 5 terms (indicated by different colors) are shown. Colors and edges indicate associated terms. Dot size indicates statistical significance as indicated by the legend.

We further hypothesized that a substantial number of misregulated genes observed in the *miR-290-295*_KO cell line (Figure 3C) may be in part a result of increased TF levels. To understand the transcriptional contribution of TFAP4 in mESCs, we attempted to rescue (i.e., downregulate) its expression level in *miR-290-295*_KO using a pool of small inhibitory RNAs (siPOOL) targeting *Tfap4* mRNA (Figure S4C). We monitored TFAP4 levels 36h after transfection by WB and observed that TFAP4 was indeed expressed at near WT levels in the *miR-290-295*_KO mESCs (Figure 4C). We then sequenced the transcriptome of the siPOOL-transfected cells and assessed the DGE (Table S7), which also confirmed near-WT *Tfap4* RNA levels (Figure 4D). In addition, we observed a striking number of DGE in the *Tfap4* siPOOL samples with a large portion of them being inversely regulated (and therefore rescued back towards WT levels) compared to the initial DE in *miR-290-295*_KO mESC lines (Figure 4D). To discriminate and quantify rescued genes, we divided the log2FC observed in *miR-290-295_*KO *vs* WT by the log2FC observed in the siPOOL-transfected miR-290-295_KO mESCs and defined rescued genes to be within the range [−0.5, −2] (orange dots in Figure 4D). We observed 287 rescued genes for which TFAP4 acted in an activating and 210 for which it acted in a repressive manner. The binding motif of TFAP4 has been previously described, based on a chromatin immunoprecipitation and sequencing (ChIP-seq) experiment in another cell type (Jackstadt et al., 2013). To refine our set of rescue-identified potential TFAP4-targets, we used PWMScan (Ambrosini et al., 2018) to scan the genome for potential TFAP4 binding sites, only keeping rescued genes with a binding site upstream of the promoter (<1kb distance), which results in 121 genes (Table S7). Finally, to better understand the role of TFAP4 in mESCs gene regulation, we performed a gene ontology analysis using the ClueGO tool on the 121 genes (Bindea et al., 2009). Most of the groups identified by the analysis, including “regulation of cell growth”, “regulation of WNT signaling pathway”, “regulation of cellular response to growth factor stimulus”, “nephron development” (Figure 4E, Table S7) are in line with previous reports of TFAP4 being an important regulator involved in developmental processes (Wong et al., 2021). In conclusion, our data demonstrate a major role for TFAP4 in stem cell gene regulation and as a TF in mouse early development.

## DISCUSSION

This study presents a novel approach that enables the accurate mapping of functional miRNA interactions in a given context by integrating computational and molecular data from various methods. We combine the well-established miRNA interaction prediction model (Agarwal et al., 2015), AGO2-binding profiles (Li et al., 2020), miRNA expression data and DGE analysis of several *RNAi*_KO mESCs, revealing 707 genes to be directly and functionally regulated by miRNAs. This corresponds to 6% of expressed genes in mESCs.

Earlier studies based on computational analyses of miRNA binding site conservation had estimated far larger numbers (60%) to be regulated by miRNAs (Friedman et al., 2009). This number was further reduced to 20% by using AGO-immunoprecipitation approach (Li et al., 2020). Our data reveal that only approximately 6% of genes are directly targeted by miRNAs in mESCs. Although this percentage is astonishingly low, relative to initial estimates, it is a reflection of the rigorous integrative method employed in this study, which aims to eliminate miRNA-mRNA interactions that are not functionally relevant in a given biological context. Moreover, Tan et al. recently developed a statistical approach for the identification of miRNA targets based on correlations in miRNA and mRNA levels in 360 lymphoblastoid cell lines (LCLs). While their approach is highly orthogonal to our integrative one, they also estimated 6% of expressed genes to be regulated by miRNAs (Tan et al., 2020), suggesting that this percentage is reflective of functionally relevant interactions in various contexts.

In order to evaluate the validity of our predictions, we knocked out the most highly expressed miRNA cluster in mESCs, miR-290-295 (Figure 3B) (Marson et al., 2008). Out of the 707 miRNA target genes identified in our integrative approach, 324 were predicted to be targeted by miR-290-295. Remarkably, the deletion of the miR-290-295 cluster revealed a statistically significant upregulation for 92 of these predicted genes, while only 9 showed significant downregulation (Figure S4A, Table S6). Most miRNA interaction prediction models rely on a similar set of features for the identification of functional miRNA interactions (Schäfer and Ciaudo, 2020). In our study, we combined the use of these established features (i.e., TargetScan score) with context-specific features (e.g., transcriptome of *RNAi*_KO mutants). The novel *miR-290-295*_KO cell lines allowed us to rank the contribution of the integrated features to the prediction of functional miRNA interactions. This demonstrated the importance of using several *RNAi*_KO cell lines for the functional validation of miRNA target genes, as applying one prediction model alone failed to accurately identify functional interactions. Our work thus overcomes the limitations of existing prediction models and implies that they should be combined with existing feature sets to better predict functional miRNA interactions in specific biological contexts such as mESCs.

Importantly, this study also identifies TFAP4 as a novel key transcription factor in stem cells, which is directly regulated by miRNAs. Moreover, gene ontology analysis reveals important roles for TFAP4 in mouse early development, cell growth roles as well as in the Wnt/β-catenin pathway. In humans, *Tfap4* has been previously described to regulate stemness and proliferation (Jackstadt et al., 2013; Jung et al., 2008), but especially to drive cancer malignancy (for review (Wong et al., 2021)). More recently, this TF has been predicted to be involved in a regulatory network of TFs involved in the reprogramming of mouse primary fibroblasts into induced pluripotent stem cells, which is consistent with our results (Papathanasiou et al., 2021). Thus, we hypothesize that TFAP4 might be an essential regulator of stemness and development. In addition to the strong impact of *Tfap4* misregulation on mESC gene expression, we also show that it is regulated by the miR-290-295 cluster (Figure 4), whose expression decreases after the exit from pluripotency of mESCs (Greve et al., 2013). MiRNA-mediated regulation of *Tfap4* has already been observed human cells, where *Tfap4* is regulated by hsa-miR-302c. Interestingly, but perhaps unsurprisingly, hsa-miR-302c shares the seed sequence with some members of the mmu-miR-290-295 cluster (Ma et al., 2018). In fact, the observation that the human miR-302c binding site is conserved in mouse ultimately contributed to the prediction of *Tfap4* as targeted by the miR-290-295 cluster in our context and leads us to believe that this miRNA-mRNA interaction is evolutionarily conserved and that *Tfap4* might therefore be regulated by miRNAs in a similar manner during human early development.

A significant revelation that has come to light from studying this single miRNA target gene is that miRNAs have the potential to act as master regulators of gene expression in regulatory networks, e.g. early development, by repressing key modulators of gene expression such as TFs. Here, we observe the complexity of miRNA-mediated regulation and its potential role in stemness and pluripotency. We therefore emphasize the importance of integrating miRNAs and other non-coding elements of the genome into our traditional understanding of gene regulatory networks. We encourage others to apply our approach to other cell types and contexts to generate accurate context-dependent miRNA interaction maps with the goal of obtaining a deeper understanding of the intricate networks that govern gene expression and concomitantly important biological processes. In many contexts and cell types, the relevant data sets have already been generated and published and await integrative analysis, as for example in the HCT116 cell line (Chu et al., 2020; Kim et al., 2016).

### Limitations of the study

A substantial part of our integrative analysis was based on transcriptomics data, and while we showed that predicted targets are also affected at the protein level, we cannot exclude that some miRNA targets are only affected at the RNA level and therefore not detected by our approach. Nevertheless, our datasets demonstrate a strong correlation between Ribo-seq and RNA-seq in mESCs as previously described in a different cellular context (Guo et al., 2010).

Another aspect to take into account is that not all miRNAs require the full set of RNAi genes for their biogenesis (Bodak et al., 2017). Our approach is partially robust against this phenomenon, as we considered genes to be potential miRNA targets even if they are upregulated in only a subset of the *RNAi*_KO mutants (i.e., in at least two). Furthermore, based on the *RNAi*_KOs’ sRNA-seq data, we identified only very few noncanonical miRNAs in mESCs and thus believe this manner is of low relevance in our context (Table S2).

Finally, as previously noted, the loss of miRNAs can lead to a cascade of downstream regulation events. In some cases, this may mask the expected upregulation in *RNAi*_KO mutants, such that functional interactions are not detectable. A similar challenge is that miRNAs are able to regulate their targets in a combinatorial manner; i.e. several miRNAs can target the same mRNA at the same time (Cursons et al., 2018). This can lead to increased repression potential as shown in Figure 4B. Disentangling such combinatorial effects requires the examination of individual and combined *miRNA*_KOs. Alternatively, machine learning models (Schäfer & Ciaudo, 2020), including graph-aware deep learning models (Zhou et al., 2020) might help to overcome some of these barriers by generating a system’s level understanding of regulation networks, potentially via the integration of a variety of regulatory mechanisms.

Our rigorous approach discards many interactions that would be falsely predicted by other methods, thus leading to a high-confidence set of direct and functional miRNA interactions in mESCs. We expect these data will be useful to the scientific community and also trust that they will serve as a robust data set on which to anchor future machine learning endeavors that can be applied to many different biological systems.

## MATERIALS AND METHODS

### Mouse ESC lines

WT E14, RNAi_KO (*Dgcr8*_KO, *Drosha*_KO, *Dicer*_KO and *Ago2&1*_KO) and *miR-290-295_KO* cluster mESC lines (129/Ola background) were cultured in Dulbecco’s Modified Eagle Media (DMEM) (Sigma-Aldrich), containing 15% fetal bovine serum (FBS; Life Technologies) tested for optimal growth of mESCs, 100 U/mL LIF (Millipore), 0.1 mM 2-ß-mercaptoethanol (Life Technologies) and 1% Penicillin/Streptomycin (Sigma-Aldrich), on 0.2% gelatin-coated support in absence of feeder cells. The culture medium was changed daily. All cells were grown at 37°C in 8% CO_2_.

### CRISPR/Cas9 mediated gene knockout

The generation of *Dgcr8*_KO, *Drosha*_KO and *Dicer*_KO mESC lines was previously described (Bodak et al., 2017a; Cirera-Salinas et al., 2017). The *Ago2&1*_KO1 and KO2 cell lines as well as *miR-290-295*_KO1 and KO2, were generated using a paired CRISPR/Cas9 strategy on WT mESCs as described previously (Wettstein et al., 2016). We generated two independent clones for the *Ago2&1*_KO line (*Ago2&1*_KO1, *Ago2&1*_KO2) using two different pairs of gRNAs to delete one or more exons of the *Ago1* gene in the previously described *Ago2*_KO1 mutant mESC line (Ngondo et al., 2018). *Ago2*_KO1 mESCs were transfected with pX458-sgRNA_*Ago1*_1/2 (Addgene #73533 and #73534), and pX458-sgRNA_*Ago1*_3/4 (Addgene #73535 and #73536) plasmids (Ngondo et al., 2018). We generated two independent *miR-290-295*_KO mESC lines by transfecting WT E14 mESCs with pX458-sgRNA_*miR290-295*_3/2 for KO1 (Addgene #172711, #172710) and pX458-sgRNA_*miR290-295*_1/2 for KO2 (Addgene #172709 and #172710). After 48 hours, the GFP positive cells were single cell sorted in 96-well plates. The deletion was genotyped by PCR using primers listed in Table S8. All transfected plasmids are available in the Addgene repository. Positive clones were expanded and verified by genomic PCR and sequencing.

### Extraction of total RNA from mESCs

Total RNA from 1-10 million cells was extracted using Trizol reagent (Life technologies) following the manufacturer’s protocol (Bodak and Ciaudo, 2016). RNA was quantified using spectrophotometry on the Eppendorf Biophotometer. RNA integrity was visually controlled by running 1 µg of total RNA extract on a 1% agarose gel.

### RNA-seq

#### Tru-seq

Prior to library preparation, the quality of isolated RNA was determined with a Bioanalyzer 2100 (Agilent, Santa Clara, CA, USA). Up to 2 µg of polyA purified RNA was used for the library preparation using the TruSeq paired-end stranded RNA Library Preparation Kit (Illumina, San Diego, CA, USA) according to the manufacturer’s recommendations. The library preparation and sequencing (Illumina HiSeq 2000) were performed by the FGCZ (Functional Genomic center, Zurich). The paired-end sequencing generated about 2×60 million reads per library.

#### QuantSeq

500 ng of total RNA was used for library preparation using the QuantSeq 3’ mRNA-Seq Library Prep Kit FWD for Illumina (Lexogen) according to the manufacturer’s recommendations. Sequencing was performed by the FGCZ (Functional Genomic center, Zurich) on the Illumina NextSeq500 platform. Single-end sequencing generated at least 20 million reads per library.

### Small RNA-seq

The Illumina TruSeq Small RNA Sample Prep Kit (Illumina, San Diego, CA, USA) was used with 1 µg of total RNA for the construction of sequencing libraries by the Functional Genomic Center Zurich (Switzerland). Sequencing was performed on an Illumina Hiseq 2500 sequencer and generated between 20 and 30 millions of single reads of 50bp per library.

### Genomic DNA extraction and PCR

Genomic DNA was extracted from 5 × 10^5^ mESCs using Phenol/Choloroform/Isoamyl Alcohol (Sigma-Aldrich). Each PCR reaction was performed using 50-100 ng of genomic DNA. Genotyping PCR primer sequences are listed in Table S8.

### Quantitative Real Time PCR Analysis of miRNAs

For miRNA quantification, 1 μg total RNA was reverse transcribed using the miScript II Reverse Transcription kit (Qiagen) according to the manufacturer’s instructions (Jay and Ciaudo, 2013). After reverse transcription, cDNA products were diluted in distilled water (1:5). Quantification of expression levels was performed on a Light Cycler 480 (Roche) using 2 µl of the diluted products, the KAPA SYBR FAST qPCR kit optimized for Light Cycler 480 (KAPA Biosystems), miScript Universal Primer (Qiagen) and a primer for the targeted miRNA. Differences between samples and controls were calculated based on the 2^−ΔΔCT^ method using RNU6 control primer (Qiagen) as normalizer. Quantitative RT-PCR assays were performed in triplicate (Jay and Ciaudo, 2013). All primers are listed in Table S8.

### Western Blot Analysis

Whole cell extracts were obtained by lysing the cells in RIPA buffer (50mM Tris-HCl pH 8, 150mM NaCl, 1% IGEPAL CA-630 (w/v), 0.5% sodium deoxycholate (w/v), 0.1% sodium dodecyl sulfate (w/v) and protease inhibitors). Protein concentrations were determined by Bradford assay (Bio-Rad Laboratories). The extracts were separated on SDS-PAGE gels and transferred to polyvinylidene fluoride membranes (Sigma Aldrich). After blocking (5% milk in TBST: 50 mM Tris-Cl, pH 7.5. 150 mM NaCl, 0.1% Tween20), membranes were incubated with primary antibodies diluted in blocking solution overnight at 4°C. Membranes were incubated with one of the following antibodies overnight: DGCR8 antibody 1:2000 (Proteintech; 10996-1-AP), DROSHA antibody 1:2000 (Cell Signaling; Cat#D28B1), DICER antibody 1:2000 (Sigma Aldrich; SAB4200087), AGO1 antibody 1:1500 (Cell Signaling; Cat#5053), AGO2 antibody 1:1500 (Cell Signaling; Cat#2897S), TFAP4 antibody 1:1000 (ab223771; Abcam), TUBULIN antibody 1:10000 (Sigma Aldrich; Cat#T6119). For secondary antibody incubation, the anti-rabbit or anti-mouse IgG HRP-linked antibody (Cell Signaling Technology) was diluted to 1:10000. Immunoblots were developed using the SuperSignal West Femto Maximum Sensitivity Substrate (Invitrogen) and imaged using the ChemiDoc MP imaging system (Bio-Rad Laboratories). TUBULIN levels or the Coomassie brilliant blue staining of the membrane are used as loading controls.

ImageJ was used to quantify band intensity for each sample, which was then normalized to the Coomassie loading control. Band intensities in the mutants were then represented as a fold-change relative to the wild type sample. Three biological replicates were used to perform the quantification.

### Transfection of mESCs with miRNA mimics

100,000 mESCs (*miR-290-295* KO2 cell line) were seeded 24 hours prior to transfection. Cells were transfected with miRNA mimics (Horizon, PerkinElmer) for miR-291a-5p (20 nM final concentration), miR-291a-3p (20 nM final concentration), or a combination of both (10 nM final concentration, each) using the Lipofectamine RNAiMax transfection reagent (Invitrogen), according to the manufacturer’s protocol. A negative control miRNA mimic was also used (Table S8). Media was changed 16 hours after transfection and cells were harvested 36 hours after transfection for protein extraction.

### mESC transfection with siPOOLs

200,000 mESCs (*miR-290-295* KO2 cell line, generated in this study) were seeded 24 hours prior to transfection. Cells were transfected with siPOOLs (siTOOLs Biotech) against *Tfap4* or a negative control siPOOL at a final concentration of 5 nM using the Lipofectamine RNAiMax transfection reagent (Invitrogen), according to the manufacturer’s protocol. Media was changed after 24 hours of transfection and cells were harvested after 36 hours of transfection using 0.05% Trypsin for further processing (RNA and protein extraction).

### Proteome analysis by SWATH-MS

The MS data acquisition (SWATH-MS and DDA mode) was performed on TripleTOF 5600 mass spectrometer equipped with a NanoSpray III source and operated by Analyst TF 1.5.1 software (AB Sciex). The samples were injected onto a C18 nanocolumn packed in-house directly in a fused silica PicoTip emitter (New Objective, Woburn, MA, USA) with 3-μm 200 Å Magic C18 AQ resin (Michrom BioResources, Auburn, CA, USA) and reverse phase peptide separation was performed on a NanoLC-Ultra 2D Plus system (Eksigent–AB Sciex, Dublin, CA, USA). The total acquired data were analyzed using a pipeline configured on the euler-Portal platform at ETH Zurich.

#### Sample preparation and protein digestion

The five distinct mESC lines (i.e., WT, *Dgcr8*_KO, *Drosha*_KO, *Dicer*_KO, and *Ago2&1*_KO) were prepared in biological duplicates (e.g., two independent CRISPR/Cas9 mutants), totaling 10 distinct samples for proteomic analysis. Corresponding cells from each 10 cm plate, were washed and scraped with ice-cold phosphate-buffered saline (PBS 1X). Then, their pellets (~ 5*10^6^ cells) collected by centrifugation at 1000 rpm, were frozen in liquid nitrogen and left at −80 °C. The cell pellets were lysed on ice using a lysis buffer containing 8 M urea (EuroBio), 50 mM NH4HCO3 (Sigma-Aldrich), and complete protease inhibitor cocktail (Roche). The mixture was sonicated at 4 °C for 5 min using a VialTweeter device (Hielscher-Ultrasound Technology) at the highest setting and centrifuged at 2500 L per sample was used for protein digestion, prior to which all samples were reduced by 5 mM tris(carboxyethyl)phosphine (Sigma-Aldrich), and alkylated by 30 mM iodoacetamide (Sigma-Aldrich). The samples, adjusted to 1.5 M UREA, were digested with sequencing-grade porcine trypsin (Promega) at a 1:50 protease/protein ratio overnight at 37 °C in 100 mM NH4HCO3 (Sigma-Aldrich). The next day, the peptide digests were purified on MicroSpin Column SilicaC18 (5-60 µg capacity, Nest Group Inc., Southborough, MA), and solubilized in 50 μ acetonitrile (ACN). The final peptide amount was determined using Nanodrop ND-1000 (Thermo Scientific), and the samples adjusted to 1 µg/µL of peptide concentration. Prior to MS injection, an aliquot of retention time calibration peptides from an iRT-Kit (RT-kit WR, Biognosys) was spiked into each sample at a 1:20 (v/v) ratio to correct relative retention times between acquisitions, and each sample injected into the duplicates (i.e., technical replicates).

#### SWATH assay library generation

The samples were recorded in data-dependent acquisition (DDA) mode to generate a mouse SWATH assay library, which is used for targeted data extraction from SWATH-MS recorded data. 15 mESC samples recorded in DDA mode were combined with 65 available DDA files originating from fractionated mouse liver peptide digest to create a common mouse assay library. The nanoLC gradient used for all acquired DDA data was linear from 2 to 35% of buffer B (i.e., 0.1% formic acid in ACN) over 120 min at a 300 nl/min flow rate. Electrospray ionization was performed in positive polarity at 2.6 kV, and assisted pneumatically by nitrogen (20 psi). Mass spectra (MS) and tandem mass spectra (MS/MS) were recorded in “high-sensitivity” mode over a mass/charge (*m/z*) range of 50 to 2000 with a resolving power of 30,000 (full width at half maximum [FWHM]). DDA selection of the precursor ions in a survey scan of 250 ms was as follows: the 20 most intense ions (threshold of 50 counts) corresponding to 20 MS/MS-dependent acquisitions of 50 ms each, charge state from 2 to 5, isotope exclusion of 4u, and precursor dynamic exclusion of 8 s leading to a maximum total MS duty cycle of 1.15 s. External mass calibration was performed by injecting a 100-fmol solution of β-galactosidase tryptic. Raw data files (.wiff) were centroided, and converted into mzXML as a final format using openMS.

The converted data files were searched in parallel using the search engines X! TANDEM Jackhammer TPP (2013.06.15.1 - LabKey, Insilicos, ISB) and Comet (version “2016.01 rev. 3”) against the ex_sp 10090.fasta database (reviewed canonical Swiss-Prot mouse proteome database, released 2017.12.01) appended with common contaminants and reversed sequence decoys (Elias and Gygi, 2007) and iRT peptide sequence. The search parameters were conducted using Trypsin digestion and allowing 2 missed cleavages. Included were ‘Carbamidomethyl (C)’ as static and ‘Oxidation (M)’ as variable modifications. The mass tolerances were set to 50 ppm for precursor-ions and 0.1 Da for fragment-ions. The identified peptides were processed and analyzed through the Trans-Proteomic Pipeline (TPP v4.7 POLAR VORTEX rev 0, Build 201403121010) using PeptideProphet (Keller et al., 2002), iProphet (Shteynberg et al., 2011), and ProteinProphet scoring. Spectral counts and peptides for ProteinProphet were filtered at FDR of 0.009158 mayu-protFDR (=0.998094 iprob). The raw spectral libraries were generated from all valid peptide spectra through automated library generation workflow on the euler-Portal platform as described earlier (Schubert et al., 2015). The final generated spectral library contained high quality MS assays for 37988 tryptic peptides from 4107 mouse proteins.

#### SWATH-MS measurement and data analysis

Reverse phase peptide separation during SWATH-MS acquisition was performed with linear nanoLC gradient from 2 to 35% of buffer B (0.1% formic acid in ACN) over 60 min at a 300 nl/min flow rate. Quadrupole settings in SWATH acquisition method were optimized for the selection of 64 variable wide precursor ion selection windows as described earlier (Röst et al., 2014). An accumulation time of 50 ms was used for 64 fragment-ion scans operating in high-sensitivity mode. At the beginning of each SWATH-MS cycle, a TOF MS scan (precursor scan) was also acquired for 250 ms at high resolution mode, resulting in a total cycle time of 3.45 s. The swaths overlapped by 1 m/z, thus covering a range of 50-2000 m/z. The collision energy for each window was determined according to the calculation for a charge 2+ ion centered upon the window with a spread of 15. Raw SWATH data files were converted into the mzXML format using ProteoWizard (version 3.0.3316) (Chambers et al., 2012), and data analysis performed using the OpenSWATH tool (Röst et al., 2014) integrated in the euler-Portal workflow. The OpenSWATH workflow input files consisted of the mzXML files from the SWATH acquired data, the TraML assay library file created above, and the TraML file for iRT peptides. SWATH data were extracted with 50 ppm around the expected mass of the fragment ions and with an extraction window of +/-300 sec around the expected retention time after performing iRT peptide alignment. The runs were subsequently aligned with a target FDR of 0.01 and a maximal FDR of 0.1 for aligned features. In the absence of a confidently identified feature, the peptide and protein intensities were obtained by integration of the respective background signal at the expected peptide retention time. The recorded feature intensities after OpenSWATH identification were filtered through R/Bioconductor package SWATH2stats (Blattmann et al., 2016) to reduce the size of the output data and remove low-quality features. The filtered fragment intensities were introduced into the R/Bioconductor package MSstats (version MSstats.daily 2.3.5), and converted to a quantification matrix of relative protein abundances using functions of data pre-processing, quality control of MS runs, and model-based protein quantification (Choi et al., 2014). Quantification matrices were used as an input data template to perform further differential analysis by One-way ANOVA test for multiple-group comparison. A TukeyHSD post hoc test revealed significant changes across control samples (WT) and four different cell line clones (i.e., *Dgcr8*_KO, *Drosha*_KO, *Dicer*_KO, and *Ago2&1*_KO). The raw counts and differential expression data are available as excel files (Table S4).

### Ribosome Profiling and data analysis

#### Ribosome Profiling sample/library preparation and sequencing

Ribosome profiling and parallel RNA-seq were performed in duplicate for WT, *Dgcr8*_KO, *Drosha*_KO, *Dicer*_KO, and *Ago2&1*_KO mESC lines, following the TruSeq Ribo Profile Kit (RPHMR12126, Illumina) with minor modifications (see below), using one 15 cm dish of confluent mESCs per replicate. Cells were briefly pretreated with cycloheximide (0.1 mg/mL) for 2 min at 37°C and then immediately harvested by scraping down in ice-cold PBS (supplemented with cycloheximide). The cell pellet was collected by brief centrifugation, snap-frozen in liquid nitrogen and stored at −80°C. From the cell pellets, lysates were prepared and ribosome-protected mRNA fragments were generated by RNase I digestion as previously described using 5 units of RNase I per OD260 (Castelo-Szekely et al., 2019). Of note, before RNase I digestion, mESC lysates were spiked-in with *Drosophila* S2 cell lysates prepared using the same lysate buffers (spike-in ratio 15_mESC_:1_S2_, based on OD260 measurements). After digestion, footprint-containing monosomes were purified via MicroSpin S-400 columns (GE Healthcare) and footprints were purified with miRNeasy Mini kit (217004 Qiagen). 5 µg fragmented RNA was used for ribosomal RNA removal using Ribo-Zero Gold rRNA Removal Kit (MRZG12324 Illumina) according to Illumina’s protocol for TruSeq Ribo Profile Kit (RPHMR12126, Illumina). Footprints were excised from 15% urea-polyacrylamide gels (with single strand RNA oligonucleotides of 26 nt and 34 nt as size markers for excision). Sequencing libraries were generated essentially following the Illumina TruSeq Ribo Profile protocol. cDNA fragments were separated on a 10% urea-polyacrylamide gel and gel slices between 70-80 nt were excised. The PCR-amplified libraries were size-selected on an 8% native polyacrylamide gel (footprint libraries were at ~150 bp). From the same initial extracts (containing the S2 lysate spike-in), parallel RNA-seq libraries were prepared essentially as described (PMID 30982898) and following the Illumina protocol. Briefly, after total RNA extraction using miRNeasy RNA Extraction kit (Qiagen), ribosomal RNA was depleted using Ribo-Zero Gold rRNA (Illumina), and sequencing libraries were generated from the heat-fragmented RNA as previously described (Castelo-Szekely et al., 2019). All libraries were sequenced in-house (Lausanne Genomic Technologies Facility) on a HiSeq2500 platform.

#### Ribosome Profiling Data Analysis

Initial analysis, including mapping and quantification of mRNA and footprint abundance, were performed as previously described (Castelo-Szekely et al., 2017). Briefly, purity-filtered reads were adapters and quality trimmed with Cutadapt v1.8 (Martin, 2011). Only reads with the expected read length (16 to 35 nt for the ribosome footprint and 35 to 60 nt for total RNA) were kept for further analysis. Reads were filtered out if they mapped to *Mus musculus* ribosomal RNA (rRNA) and transfer RNA (tRNA) databases (ENSEMBL v91, (Cunningham et al., 2019)) using bowtie2 v2.3.4.1 (Langmead and Salzberg, 2012). The filtered reads were aligned against *Mus musculus* transcripts database (ENSEMBL v91) using bowtie2 v2.3.4.1. Finally, remaining reads were mapped against *D. melanogaster* transcript database (ENSEMBL v78). Reads mapping to transcripts belonging to multiple gene loci were filtered out. Reads were then summarized at a gene level using an in-house script and mouse samples were then normalized by using corresponding fly spike-in read counts. Differential ribosome occupancy was defined by DESeq2 with an absolute fold change > 0.5 and FDR (adjusted p-value) < 0.05. The spike-in normalized counts and differential expression data are available as excel files (Table S5).

### sRNA-seq analysis

Reads were trimmed using Cutadapt 1.13 (Martin, 2011) with adapter TGGAATTCTCGGGTGCCAAGG and arguments “-m 14 -M 40” and aligned to the mouse genome (GRCm38 primary assembly, annotation: GENCODE vM20) using STAR 2.4.2a (Dobin et al., 2013) with arguments “--outFilterMismtachNoverLmax 0.05” to allow for 0 mismatches for reads < 20bp. Next, reads were counted using subread-featureCounts 1.5.0 (Liao et al., 2014) with arguments “-f -O -s 1 --minOverlap 17” for 3 different annotations (miRbase v21 (Griffiths-Jones et al., 2006), GENCODE vM20 and GtRNAdb2 mm10 (Chan and Lowe, 2016) to collect gene counts for the following groups of small RNAs: miRNAs, small nuclear RNAs, small nucleolar RNAs, mitochondrial tRNAs and tRNAs. To take the global reduction of small RNAs in *RNAi*_KO mESC lines into account, miRNA expression was normalized by small nuclear RNA, small nucleolar RNA, mitochondrial tRNA and tRNA expression using estimateSizeFactors from DESeq2 1.18.1 (Love et al., 2014). Read counts normalized in this way are referred to as *CPM-like* in this manuscript.

### RNA-seq analysis

Raw and normalized read counts for both genes and transcripts were computed by the RNA-seq pipeline from snakePipes 2.3.1 (Bhardwaj et al., 2019) using the ENSEMBL GRCm38.98 primary assembly and annotation. The following command line arguments were passed “--trim --trimmer trimgalore --trimmerOptions ‘--illumina --paired’ --mode ‘alignment,alignment-free,deepTools_qc -- fastqc”. The RNA-seq pipeline was further run for each *RNAi*_KO mESC line with an according sample sheet and the “--sampleSheet” option to perform differential gene expression (DGE) analysis. Briefly, the pipeline employs TrimGalore/Cutadapt (Martin, 2011), STAR (Dobin et al., 2013), featureCounts (Liao et al., 2014) and DESeq2 (Love et al., 2014) to produce read counts and DGE data on a per-gene basis. Salmon (Patro, 2017) was employed to derive per-transcript expression.

### QuantSeq analysis

In accordance with the QuantSeq manual, first, adapters were trimmed using TrimGalore with arguments “--stringency 3 --illumina” then the polyA tail was trimmed using TrimGalore with argument “--polyA”. Next, snakePipes (Bhardwaj et al., 2019) RNA-seq pipeline was run with the arguments “--libraryType 1 --featureCountOptions ‘--primary’ --mode alignment,alignment-free,deepTools_qc”. The RNA-seq pipeline was further run for *miR-290-295*_KO mESC line with an according sample sheet and the additional “--sampleSheet” option to perform DGE analysis.

### Integration of multi-OMICs miRNA interaction data

#### Retrieval and preparation of TargetScan score

Conserved and nonconserved site context score tables were downloaded from TargetScan mouse 7.2 (Agarwal et al., 2015) and concatenated. The context++ score from these data is referred to as *TargetScan score* in this manuscript.

#### AGO2 binding data analysis

AGO2 binding peaks were downloaded from GEO (GSE139345) as provided by (Li et al., 2020) and miRNA seed matches (7merA1, 7merm8 and 8mer; 6mers were discarded) in peaks were identified for all mouse miRNAs (miRBase v21 (Griffiths-Jones et al., 2006)). Seed matches were then mapped to gene regions to further associate them with a gene ID (if applicable) and the region type (5’UTR, CDS or 3’UTR).

#### Integration of AGO2 binding, TargetScan, miRNA expression and gene upregulation in RNAi KO mESC lines

AGO2 binding data and TargetScan scores were preprocessed as explained above and reduced such that there was only a single entry for each (gene, region_type, miRNA, seed_match_type)-tuple. In the rare cases of duplicates, the associated scores (peak size for AGO2 binding data and context++ score for TargetScan interactions) were computed as the exponentially decaying weighted sum (e.g. 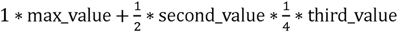). Next, prepared TargetScan scores were joined on the set of unique keys (gene, region_type, miRNA, seed_match_type) such that data entries without a match in one of the datasets were kept and missing data fields were set to 0 (as in an outer-join operation). The result was stored as integrated, unfiltered set of interactions.

#### Filtering and scoring of integrated interaction data

Filtering was performed in three steps. First, interactions with an AGO2-binding value of 0 were deleted. Next, interactions where the corresponding miRNA was expressed with less than 10 CPM-like in WT mESCs were deleted. Finally, interactions were deleted where the corresponding mRNA was not in the set of upregulated genes. This set was defined as follows: log2(fold-change) should be higher than −0.4 in all, and higher than 0.5 (with an adjusted p-value < 0.1) in at least two *RNAi*_KO mutants. Filtering for TargetScan’s context++ score (< −0.15) was only performed for comparison and not for downstream analysis.

To allow for a confidence ranking of interactions, the mean of the four feature scores WT miRNA expression, AGO2 binding, TargetScan score and mutant upregulation was used as Interaction score. The four scores were produced by scaling to [0, 1] after applying the log2 to the miRNA expression and AGO2 binding peak enrichment. The mutant upregulation was computed as count of mutants with stat. sign. upregulation (adj. p-value < 0.1, log2(fold-change) > 0.5).

To allow for a ranking of genes, such that high confidence targets of miRNAs are ranked highest, interaction scores were grouped and combined on a per-gene basis in the following manner: the mutant upregulation score (which is logically the same for all interactions of the same gene) was added to the geometric mean of the maximum and of the sum of the three summed miRNA-associated features. Here, the rationale was to rank genes higher if they were subject to larger numbers of interactions, however, to avoid high ranking of genes with a large number of interactions with low scores, the geometric mean dampened their score while favoring genes with large numbers of interactions *and* interactions with high scores.

### Gene ontology analysis of *Tfap4* targets

Potential *Tfap4*-targets (as determined from their degree of rescue in the siPOOL-treated cells) were further filtered for TFAP4-binding sites in the genome. The PWMScan website (Ambrosini et al., 2018) was used with default parameters. The GRCm38/mm10 genome was selected and scanned for human CIS-BP TFAP4 binding motifs. The resulting bed file was downloaded and used to select those genes that had a binding motif closely upstream to their transcription start site (<1kb distance). Gene ontology analysis was performed for these genes using ClueGO (Bindea et al., 2009) with the following options: Network specificity was set to medium-1, GO term fusion was enabled, only pathways/terms with pV < 0.05 were shown and terms from WikiPathways, KEGG, GO Biological Processes, GO Cellular Components and GO Molecular Function from 2021/05/13 were used.

### Custom data analyses, visualizations

Data analyses and visualizations were realized as described in the last sections using bash and python scripting, organized in a Snakemake pipeline (Mölder et al., 2021). PCA analysis was performed using scikit-learn (Pedregosa et al., 2011).

## Acknowledgments

We would like to thank the members of the Ciaudo lab and Dr. Tobias Beyer for fruitful discussions and the critical reading of this manuscript. We are grateful to Bulak Arpat for help at the beginning of computational analyses of the Ribo-seq datasets. This work was supported by the Swiss National Science Foundation (grants 31003A_173120 and 310030_196861) to C.C. C.C, H.W and D.G were supported by the NCCR RNA and Disease. We also want to thank the Functional Genomics Center Zurich (FGCZ) for their support with the preparation of RNA-seq libraries and sequencing.

## Author Contributions

Conceptualization, MS, DS and CC; laboratory experiments, MS, AN, MB, HW, PS, RPN; Full proteome: TS; Ribo-seq libraries preparation, AL; computational analysis, MS and DS; writing—original draft preparation, MS, AN and CC; writing—review and editing, CC; expertise and editing, RA and DG; visualization, MS, AN, MB and CC.; supervision, CC; funding acquisition, CC. All authors have read and agreed to the published version of the manuscript.

---

**Supplementary Figure 1:**
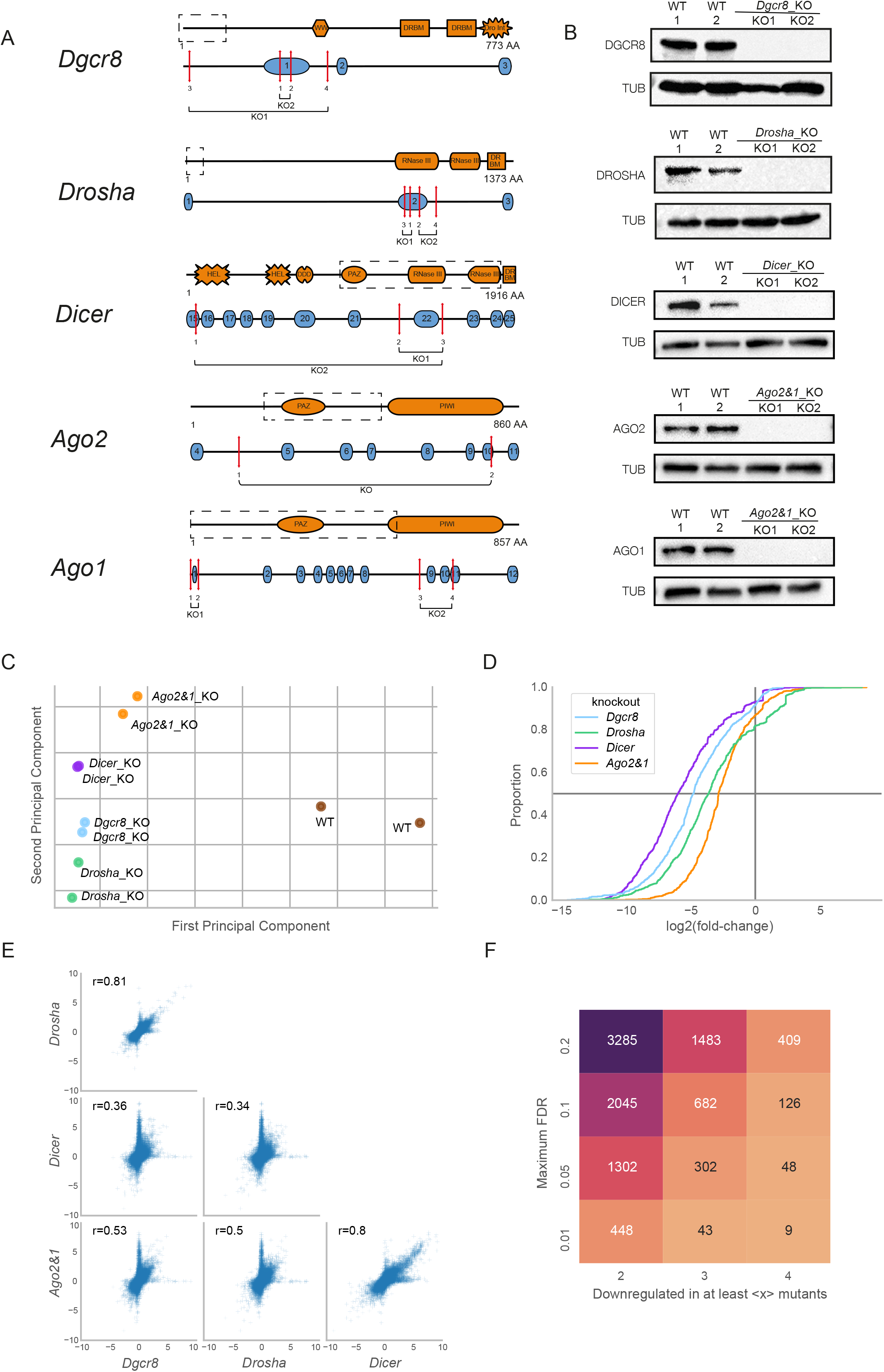
Characterization of *RNAi*_KO mESC lines. Related to Figure 1. (A) Schematic representation of paired CRISPR-Cas9 KO strategies for the generation of *RNAi*_KO mESC lines. Red arrows indicate loci targeted by sgRNAs. (B) Immunoblot analysis of RNAi proteins in WT and *RNAi*_KO mESC lines. TUBULIN was used as a loading control. Representative blot of three independent experiments is shown. (C) PCA plot of miRNA expression as measured by sRNA-seq in *RNAi*_KO and WT samples. Biological replicates are indicated with the same color. (D) Cumulative distribution function of differential miRNA expression in *RNAi*_KO versus WT mESCs as measured by sRNA-seq. Values smaller than zero indicate reduced expression as compared to WT. (E) Pairwise scatterplot matrix comparing DEG (log2FoldChanges) in *RNAi*_KO mESCs. Each small cross represents a single gene and the correlation between each pair of samples is represented by Pearson correlation in the top-left corner of each sub-plot. (F) Heatmap indicating the number of genes commonly downregulated across mutants for two different selection dimensions: Y-axis denotes maximum false discovery rate (FDR) to consider a gene as downregulated. Matrix fields correspond to the number of downregulated genes, identified in at least 2, 3 or 4 mutants, as denoted by x-axis.

**Supplementary Figure 2:**
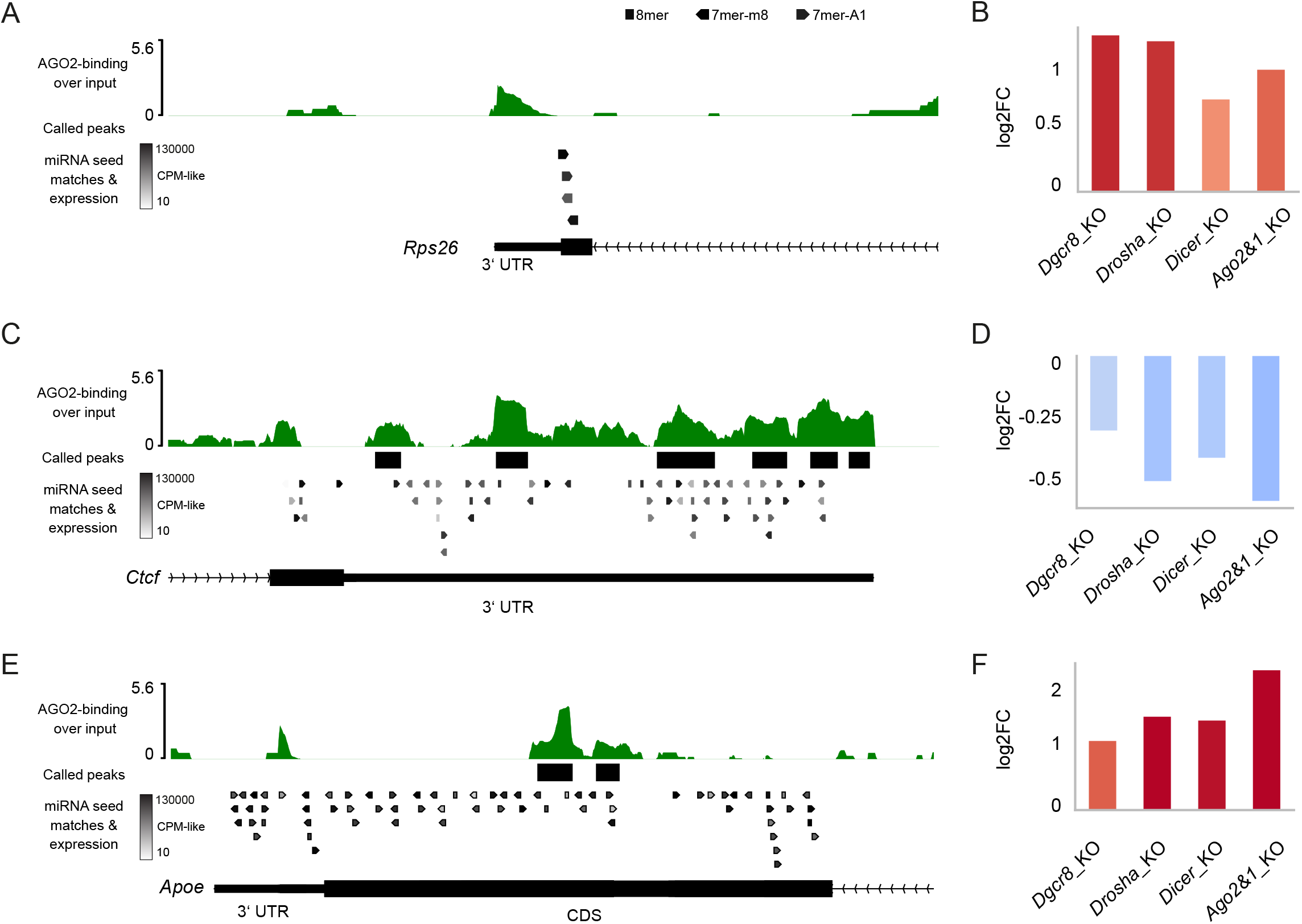
Examples of integrated data for genes with contradictory signals. Related to Figure 2. (A, B) No statistically significant AGO2-binding in 3’UTR of the *Rps26* gene and no 8mer binding sites in 3’UTR, yet with observed upregulation in all *RNAi*_KO mESC lines. (C, D) AGO2-binding in 3’UTR of the *Ctcf* gene with multiple potent binding sites, yet without observed upregulation in *RNAi*_KO mESC lines. (E, F) No statistically significant AGO2-binding in 3’ UTR of the *Apoe* gene, yet statistically significant binding to the CDS at a strongly expressed 8mer binding site along with a consistent upregulation in all four mutants. For a more detailed subplot description, refer to the figure legend of Figures 2B, C, D, E.

**Supplementary Figure 3:**
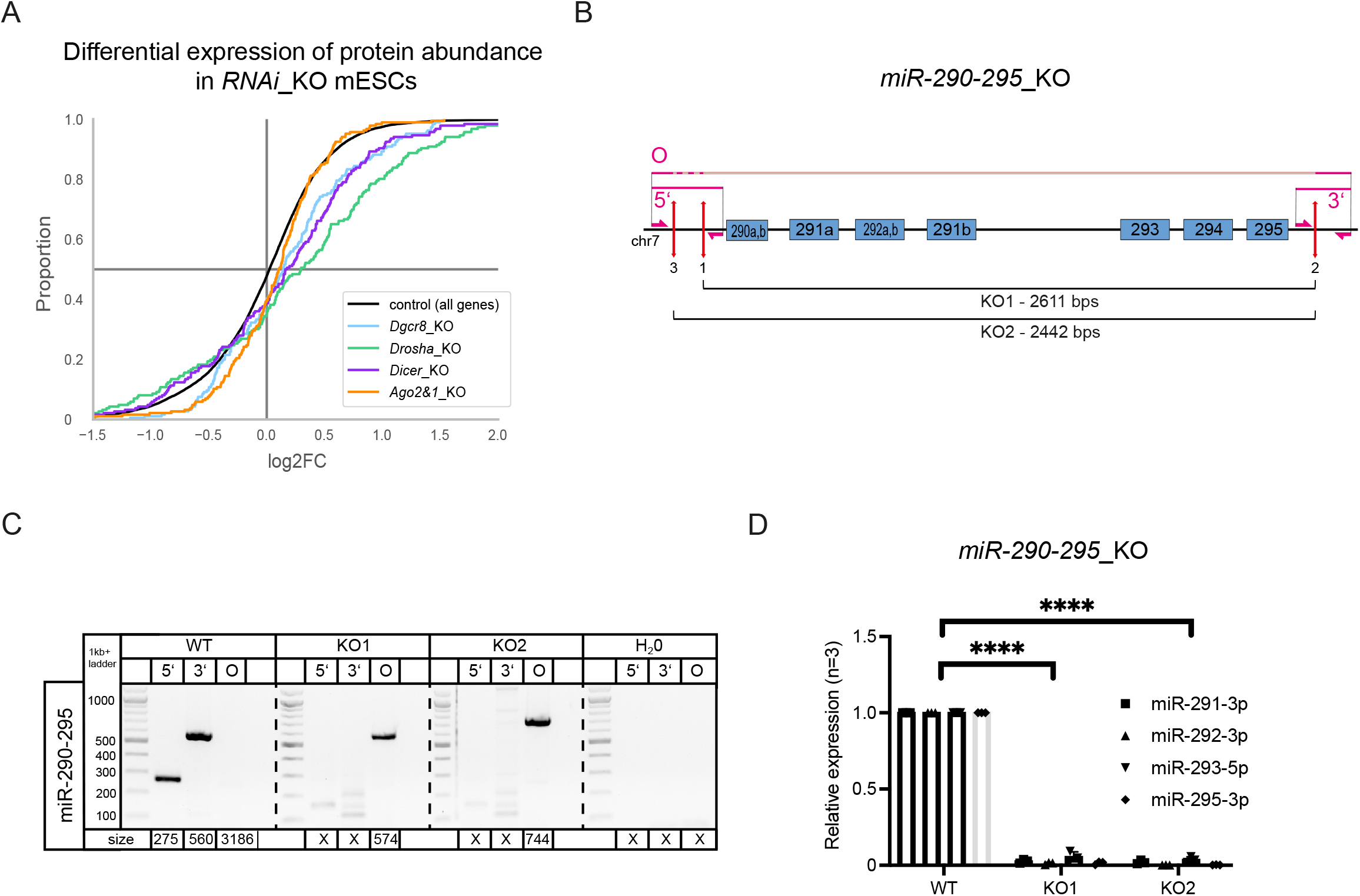
Full proteome analysis and characterization of miR-290-295_KO mESC lines. Related to Figure 3. (A) Cumulative distribution function of differential protein abundance as detected by SWAT-MS in *RNAi*_KO mESCs. Of 707 predicted miRNA target genes, 186 (26%) were detectable in the SWATH-MS data and appear in the plot. Enrichments for positive log2FoldChanges were significant for all four mutants (p<0.002 for every mutant, student’s t-test). (B) Schematic representation of the paired CRISPR/Cas9 KO approach for the generation of *miR-290-295*_KO cluster mESC lines. Genomic loci of miRNAs are indicated by blue boxes. Red lines mark sgRNA positions. Primers for screening of the genomic deletion are indicated by pink half-sided arrows and pink lines show PCR products. (C) Genomic PCR screening of *miR-290-295*_KO mESCs. Three screening primers, one around the 5’ cut site, one around the 3’ cut site and one around the entire KO region (O) were used, as annotated in Figure S3B. Expected amplicon sizes are denoted below the lanes. An *X* denotes no expected product. (D) Quantitative PCR for representative miR-290-295 members in WT and *miR-290-295_*KO mESC lines. Expression levels were normalized to RNU6. Statistical comparison was performed using 2-way ANOVA test. **** p < 0.0001.

**Supplementary Figure 4:**
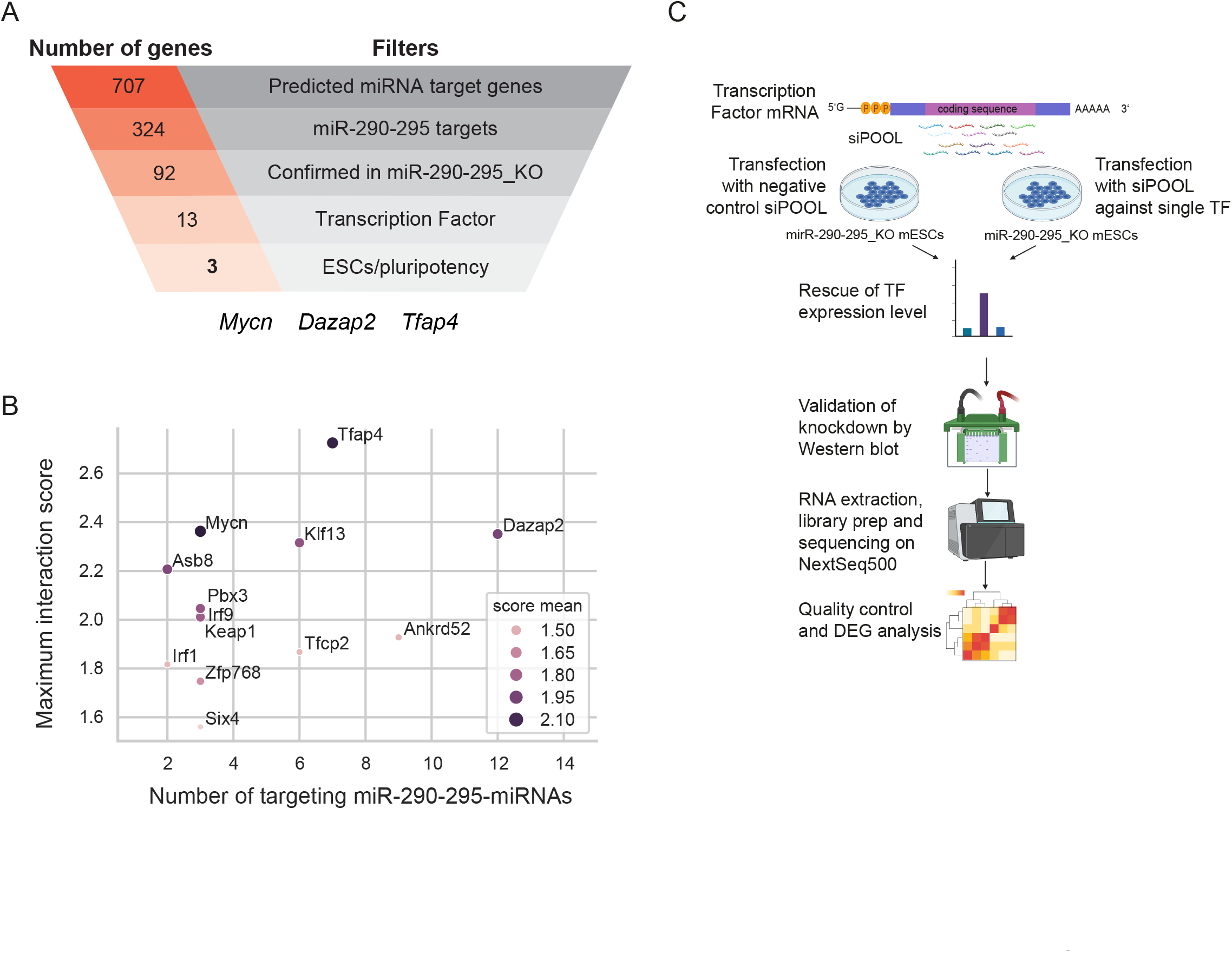
*Tfap4* is a high-confidence target of miRNAs in mESCs. Related to Figure 4. (A) Funnel analysis representing *miR-290-295* regulated transcription factors previously implicated in stem cell functions and pluripotency in the scientific literature. (B) Interaction scores from our integrative analysis (Figure 2) for all transcription factors predicted to be targeted by *miR-290-295*. Number of distinct miRNA binding sites for the miR-290-295 cluster, mean of interaction score and maximum of interaction score are shown. (C) Schematic representation describing the workflow to rescue *Tfap4* expression by treating *miR-290-295*_KO2 mESCs with pool of siRNAs (siPOOL) targeting *Tfap4*. Transfected cells were harvested and analyzed on the transcriptomics level using RNA-seq. Illustrations were extraction from BioRender.com.

**Table S1: RNA-seq TPM values and differential expression for WT and/versus RNAi_KO mESCs. Related to Figure 1.**

**TableS2: sRNA-seq (miRNAs) CPM values and differential expression for WT and/versus RNAi_KO mESCs. Related to Figures S1C, D.**

**Table S3: Filtered interactions from integrative analysis. Related to Figure 2E.**

**Table S4: Differential expression analysis for protein abundances in RNAi_KO versus WT mESCs. Related to Figure S3A.**

**Table S5: Ribo-seq differential expression analysis from *RNAi*_KO versus WT mESCs. Related to Figure 3A.**

**Table S6: QuantSeq NGS CPM values and differential expression for miRNA 290-295_KO mESCs. Related to Figure 3C.**

**Table S7: QuantSeq NGS CPM values and DE for siPOOL-knockdown of Tfap4 transcription factor. Related to Figure 4D.**

**Table S8: Primers list.**

